# Genomic analysis of early SARS-CoV-2 strains introduced in Mexico

**DOI:** 10.1101/2020.05.27.120402

**Authors:** Blanca Taboada, Joel Armando Vazquez-Perez, José Esteban Muñoz Medina, Pilar Ramos Cervantes, Marina Escalera-Zamudio, Celia Boukadida, Alejandro Sanchez-Flores, Pavel Isa, Edgar Mendieta Condado, José Arturo Martínez-Orozco, Eduardo Becerril-Vargas, Jorge Salas-Hernández, Ricardo Grande, Carolina González-Torres, Francisco Javier Gaytán-Cervantes, Gloria Vazquez, Francisco Pulido, Adnan Araiza Rodríguez, Fabiola Garcés Ayala, Cesar Raúl González Bonilla, Concepción Grajales Muñiz, Víctor Hugo Borja Aburto, Gisela Barrera Badillo, Susana López, Lucía Hernández Rivas, Rogelio Perez-Padilla, Irma López Martínez, Santiago Ávila-Ríos, Guillermo Ruiz-Palacios, José Ernesto Ramírez-González, Carlos F. Arias

**Affiliations:** Departamento de Genética del Desarrollo y Fisiología Molecular, Instituto de Biotecnología, Universidad Nacional Autónoma de México, Cuernavaca, Morelos, Mexico; Instituto Nacional de Enfermedades Respiratorias Ismael Cosío Villegas, Ciudad de México, Mexico; División de Laboratorios de Vigilancia e Investigación Epidemiológica, Instituto Mexicano del Seguro Social, Ciudad de México, México; Instituto Nacional de Ciencias Médicas y Nutrición, Ciudad de México, Mexico; Department of Zoology, Oxford University, Oxford, UK; Centro de Investigación en Enfermedades Infecciosas, Instituto Nacional de Enfermedades Respiratorias Ismael Cosío Villegas, Ciudad de México, Mexico; Unidad Universitaria de Secuenciación Masiva y Bioinformática, Instituto de Biotecnología, Universidad Nacional Autónoma de México, Cuernavaca, Morelos, Mexico; Instituto de Diagnóstico y Referencia Epidemiológicos, Dirección General de Epidemiología, Ciudad de México, Mexico; División de Desarrollo de la Investigación, Instituto Mexicano del Seguro Social, Ciudad de México, México; Coordinación de Investigación en Salud, Instituto Mexicano del Seguro Social, Ciudad de México, México; Coordinación de Control Técnico de Insumos, Instituto Mexicano del Seguro Social, Ciudad de México, México; Dirección de Prestaciones Médicas, Instituto Mexicano del Seguro Social, Ciudad de México, Mexico

**Author notes:** Corresponding authors: José Ernesto Ramírez-González, and Carlos F. Arias, JERG: Instituto de Diagnóstico y Referencia Epidemiológicos, Dirección General de Epidemiología, Ciudad de México, Mexico., CFA: Departamento de Genética del Desarrollo y Fisiología Molecular, Instituto de Biotecnología, Universidad Nacional Autónoma de México, Cuernavaca, Morelos, Mexico. Blanca Taboada, Joel Armando Vazquez-Perez, José Esteban Muñoz Medina, Pilar Ramos Cervantes, and Marina Escalera-Zamudio contributed equally to this work.

## Abstract

The COVID-19 pandemic has affected most countries in the world. Studying the evolution and transmission patterns in different countries is crucial to implement effective strategies for disease control and prevention. In this work, we present the full genome sequence for 17 SARS-CoV-2 isolates corresponding to the earliest sampled cases in Mexico. Global and local phylogenomics, coupled with mutational analysis, consistently revealed that these viral sequences are distributed within 2 known lineages, the SARS-CoV-2 lineage A/G, containing mostly sequences from North America, and the lineage B/S containing mainly sequences from Europe. Based on the exposure history of the cases and on the phylogenomic analysis, we characterized fourteen independent introduction events. Additionally, three cases with no travel history were identified. We found evidence that two of these cases represent local transmission cases occurring in Mexico during mid-March 2020, denoting the earliest events described in the country. Within this Mexican cluster, we also identified an H49Y amino acid change in the spike protein. This mutation is a homoplasy occurring independently through time and space, and may function as a molecular marker to follow on any further spread of these viral variants throughout the country. Our results depict the general picture of the SARS-CoV-2 variants introduced at the beginning of the outbreak in Mexico, setting the foundation for future surveillance efforts.

This work is the result of the collaboration of five institutions into one research consortium: three public health institutes and two universities. From the beginning of this work, it was agreed that the experimental leader of each institution would share the first authorship. Those were the criteria followed to assign first co-first authorship in this manuscript. The order of the other authors was randomly assigned.

**IMPORTANCE:** Understanding the introduction, spread and establishment of SARS-CoV-2 within distinct human populations is crucial to implement effective control strategies as well as the evolution of the pandemics. In this work, we describe that the initial virus strains introduced in Mexico came from Europe and the United States and the virus was circulating locally in the country as early as mid-March. We also found evidence for early local transmission of strains having the mutation H49Y in the Spike protein, that could be further used as a molecular marker to follow viral spread within the country and the region.

## INTRODUCTION

The 2019 coronavirus disease (COVID-19), declared a pandemic by the WHO on March 11^th^, 2020^1^, is caused by a novel betacoronavirus known as the severe acute respiratory syndrome coronavirus 2 (SARS-CoV-2), detected in December of 2019 in the province of Wuhan in China^1^. This is the third outbreak related to zoonotic betacoronaviruses known to occur in humans in the last two decades, after SARS (severe acute respiratory syndrome) in 2002 and MERS (Middle East respiratory syndrome) in 2012. After its emergence in China, SARS-CoV-2 spread initially to other parts of the world by people with a travel history to China, but gradually shifted to local transmissions^2^. Viral spread was first detected in Thailand, South Korea and Japan, and by the second half of January, the first positive cases appeared in the USA and Europe (France, Italy and Spain). The current SARS-CoV-2 genome analysis in the Nextstrain site^3^, points out that viral transmission is now mainly community-driven^4,5^.

In many countries, despite diagnostic efforts and initial control strategies, SARS-CoV-2 spread went on undetected until a critical number of cases requiring hospitalization and intensive care was reached, alerting the authorities in charge. As for May 4^th^ 2020, SARS-CoV-2 has infected more than 3,578,000 people and caused around 251,000 deaths worldwide^3^. In Mexico, the first case of SARS-CoV-2 was detected on February 27^th^ 2020, corresponding to a person who travelled back to Mexico from Italy, and who was in direct contact with a confirmed SARS-CoV-2 case. Soon after, additional cases were detected among travelers that returned from the USA and Europe, increasing every day. By May 4^th^, there were over 23,400 confirmed cases and 2,150 deaths within the country, indicating local transmission^6^. Understanding the introduction, spread and establishment of SARS-CoV-2 within distinct human populations is crucial to implement effective control strategies. In this work, we studied the early introduction dynamics of the first SARS-CoV-2 cases in Mexico. For this, we used a whole genome sequencing and phylogenomic approach. We obtained 17 full viral genome sequences, including the first case detected and sampled within the country. Phylogenomic placement showed that these viruses belong to the A2/G and B/S lineages, two of the three circulating viral lineages reported so far. Our analysis also confirms that there have been multiple independent introduction events in Mexico from travelers abroad. We also found evidence for early local transmission of strains having the mutation H49Y in the Spike protein, that could be further used as a molecular marker to follow viral spread within the country.

## METHODS

### Ethical statement

All clinical samples were processed at the “Instituto de Diagnóstico y Referencia Epidemiológicos” (InDRE), following official procedures^7^. All samples used for this work are considered part of the national response to COVID-19, and the data collected is directly related to disease control.

### Sample collection and diagnostics

All samples used in this study were collected under the Mexican Official Norm NOM-024-SSA2-1994 for prevention and control of acute respiratory infections in the primary health attention, as part of the early diagnostics scheme for SARS-CoV-2 in public health laboratories and hospitals in Mexico City (Red Nacional de Laboratorios Estatales de Salud Pública, RNLSP; Instituto Nacional de Enfermedades Respiratorias, INER; Instituto Nacional de Ciencias Médicas y Nutrición Salvador Zubirán, INCMNSZ; and Instituto Mexicano del Seguro Social, IMSS). Oro- and naso-pharyngeal swabs were collected and placed in virus transport medium upon collection, following InDRE official procedures^8^. A tracheal aspirate was also obtained from one patient and it was frozen at −70°C until use. Diagnosis was done using validated protocols for SARS-CoV 2, as approved by InDRE and by the World Health Organization (WHO)^9^.

### Sample processing and Whole Genome Sequencing

All samples were prepared for RNA extraction, as described^10,11^. Briefly, centrifuged and filtered supernatants were treated with Turbo DNase and RNAse. Nucleic acids were then extracted using the PureLink™ Viral RNA/DNA Kit (ThermoFisher), following the manufacturer’s instructions and using linear acrylamide (Ambion) as RNA carrier. cDNA was synthesized using the SuperScript III Reverse Transcriptase System (ThermoFisher) and primer A (5’-GTTTCCCAGTAGGTCTCN9-3’). The second strand was generated by two rounds of synthesis with Sequenase 2.0, followed by 15 cycles of amplification using Phusion DNA polymerase using primer B (5’-GTTTCCCAGTAGGTCTC-3). Next, cDNA was purified using the DNA Clean & Concentrator Kit (Zymo Research) and used as input material for generating sequencing libraries, following the Nextera XT DNA Library Preparation Kit^12^ (Illumina). Finally, all samples were sequenced on the Illumina NextSeq 500 platform using a 150 cycle High Output Kit v2.5 to obtain paired end reads of 75 base pairs. Sequencing yields are reported in SI Table 1.

**Table 1.** List of viral genomes derived from Mexican samples.

| Sample ID | Virus name | Accession<br>GISAID ID | Location<br>Country City | Age | Sex M/F | EH Place | Collection<br>date | Arrival to<br>Mexico | Port of<br>Entry |
| --- | --- | --- | --- | --- | --- | --- | --- | --- | --- |
| 2 | Mexico/CDMX-INER_01 | EPI_ISL_424345 | Mexico_CDMX | 42 | M | EH_Spain | 12/03/20 | 12/03/20 | Mexico City |
| 5 | Mexico/CDMX-INER_02 | EPI_ISL_424348 | Mexico_CDMX | 55 | F | EH_Spain_France | 13/03/20 | 12/03/20 | Mexico City |
| 6 | Mexico/CDMX-INER_03 | EPI_ISL_424625 | Mexico_CDMX | 29 | M | EH_Spain | 13/03/20 | 11/03/20 | Mexico City |
| 7 | Mexico/CDMX-INER_04 | EPI_ISL_424626 | Mexico_CDMX | 25 | F | EH_Egypt_Barcelona_Spain | 15/03/20 | 15/03/20 | Mexico City |
| 8 | Mexico/CDMX-INER_05 | EPI_ISL_424627 | Mexico_CDMX | 38 | M | EH_None | 15/03/20 | N/A | N/A |
| 13 | Mexico/Chihuahua-IMSS_01 | EPI_ISL_424731 | Mexico_Chihuahua | 21 | M | EH_UK_France_USA | 13/03/20 | 12/03/20 | Mexico City |
| 16 | Mexico/Chiapas-InDRE_02 | EPI_ISL_424666 | Mexico_Chiapas | 18 | F | EH_Italy | 29/02/20 | 25/02/20 | Mexico City |
| 17 | Mexico/EdoMex-InDRE_03 | EPI_ISL_424667 | Mexico_Edomex | 71 | M | EH_Italy | 04/03/20 | 21/02/20 | Mexico City |
| 19 | Mexico/Queretaro-InDRE_04 | EPI_ISL_424670 | Mexico_Queretaro | 43 | M | EH_Spain_Holand_USA | 10/03/20 | 06/03/20 | Mexico City |
| 22 | Mexico/Puebla-InDRE_05 | EPI_ISL_424672 | Mexico_Puebla | 31 | M | EH_Spain_France | 11/03/20 | 09/03/20 | Mexico City |
| 24 | Mexico/CDMX-InDRE_06 | EPI_ISL_424673 | Mexico_CDMX | 63 | F | EH_Denver_USA | 12/03/20 | 06/03/20 | Mexico City |
| 27 | Mexico/CDMX-INCMNSZ_01 | EPI_ISL_426361 | México_CDMX | 70 | F | EH_Vail_USA | 12/03/20 | 08/03/20 | Mexico City |
| 28 | Mexico/CDMX-INCMNSZ_02 | EPI_ISL_426362 | México_CDMX | 70 | M | EH_Vail_USA | 12/03/20 | 08/03/20 | Mexico City |
| 30 | Mexico/CDMX-INCMNSZ_03 | EPI_ISL_426363 | México_CDMX | 59 | M | EH_Madrid_Spain | 08/03/20 | 12/03/20 | Mexico City |
| 31 | Mexico/CDMX-INCMNSZ_04 | EPI_ISL_426364 | México_CDMX | 71 | M | EH_Vail_USA_Germany | 10/03/20 | 08/03/20 | Mexico City |
| 32 | Mexico/CDMX-INCMNSZ_05 | EPI_ISL_426365 | México_CDMX | 32 | M | EH_None | 12/03/20 | N/A | N/A |
| 33 | Mexico/CDMX-InDRE_01 | EPI_ISL_412972 | México_CDMX | 35 | M | EH_Italy | 27/02/20 | N/A | Mexico City |

### Bioinformatic analysis

#### Data quality control and processing

Read quality control was carried out using FAST-QC^13^ under the default parameters. Adapter sequences and low-quality bases were removed using Fastp v0.19^14^. Low complexity reads, those with a length shorter than 40 bases, and duplicates were excluded using CD-HIT-DUP v.4.6.8^15^. Off-target reads were then filtered out using Bowtie2 v2.3.4.3 with the default parameters against the human genome version GRCh38.p13, and the SILVA database^16^ as a reference to filter out human DNA and ribosomal sequences.

#### Viral genome assembly

The reads obtained were used as input to assemble viral genomes using the Wuhan-Hu-1 reference genome sequence (MN908947). For this, the reads obtained for each sample were mapped against the reference using Bowtie2 v2.3.4.3. Aligned reads were then used for *de novo* assembly using SPAdes v^17^. Consensus genome sequences were generated using the majority threshold criterion. Only sequences with a coverage above 80% and a mean depth of ≥8X were considered for the analyses (SI Table 1).

### Phylogenetic analyses

#### Data collation

From 4698 complete SARS-CoV-2 genomes deposited in the Global Initiative on Sharing All Influenza Data (GISAID) platform on the morning of April 7^th^ 2020, a total of 3014 sequences genomes (>29,000 nt and only high coverage) were downloaded to generate a local database. As the collection dates of the Mexican samples range from late February to March, we filtered sequences collected between 02/01/2020 to 03/31/2020 for our database (Table 1). From these, unique sequences were extracted and those identical were collapsed, leaving 2633. We then included the 17 consensus viral genomes determined in this study and the Wuhan-Hu-1 reference genome sequence, yielding a total of 2651 sequences. We aligned the whole genome nt dataset using MUSCLE v3.8^18^ under default parameters, and then used the *getorfs* script from the EMBOSS suite^19^ to extract complete ORFs (Open Reading Frames) above 300 nt (Orf1a, Orf1b, Spike, M, Orf3a, Orf7a, Orf8 and N), that were then individually re-aligned as described^18^. Finally, to exclude UTRs and non-coding intergenic regions from the phylogenomic analyses, individual ORFs were concatenated to generate an additional 28,320 nt-long whole genome (WG) alignment.

#### Data subsampling and tree inference

The individual and concatenated alignments were then reversed to nucleotides and used for estimating maximum likelihood (ML) trees using RAxML v8^20^ under the following parameters: -T 2 -f a -x 390 -m GTRGAMMA -p 580 -N 100. All trees were rooted on Wuhan-Hu-1 reference genome sequence^21^. Given that the SARS-CoV-2 virus shows a low degree of genetic variation^22^, clade definition must be based on consensus branching patterns within different trees and by shared nucleotide substitution patterns^23^, in addition to bootstrap support values. Based on these criteria, the position of the Mexican sequences was determined within the whole genome (WG) and individual ORF1a, ORF1b and S trees (SI Table 2), and was then confirmed on the global phylogeny available in Nextstrain^24^.

**Table 2.**
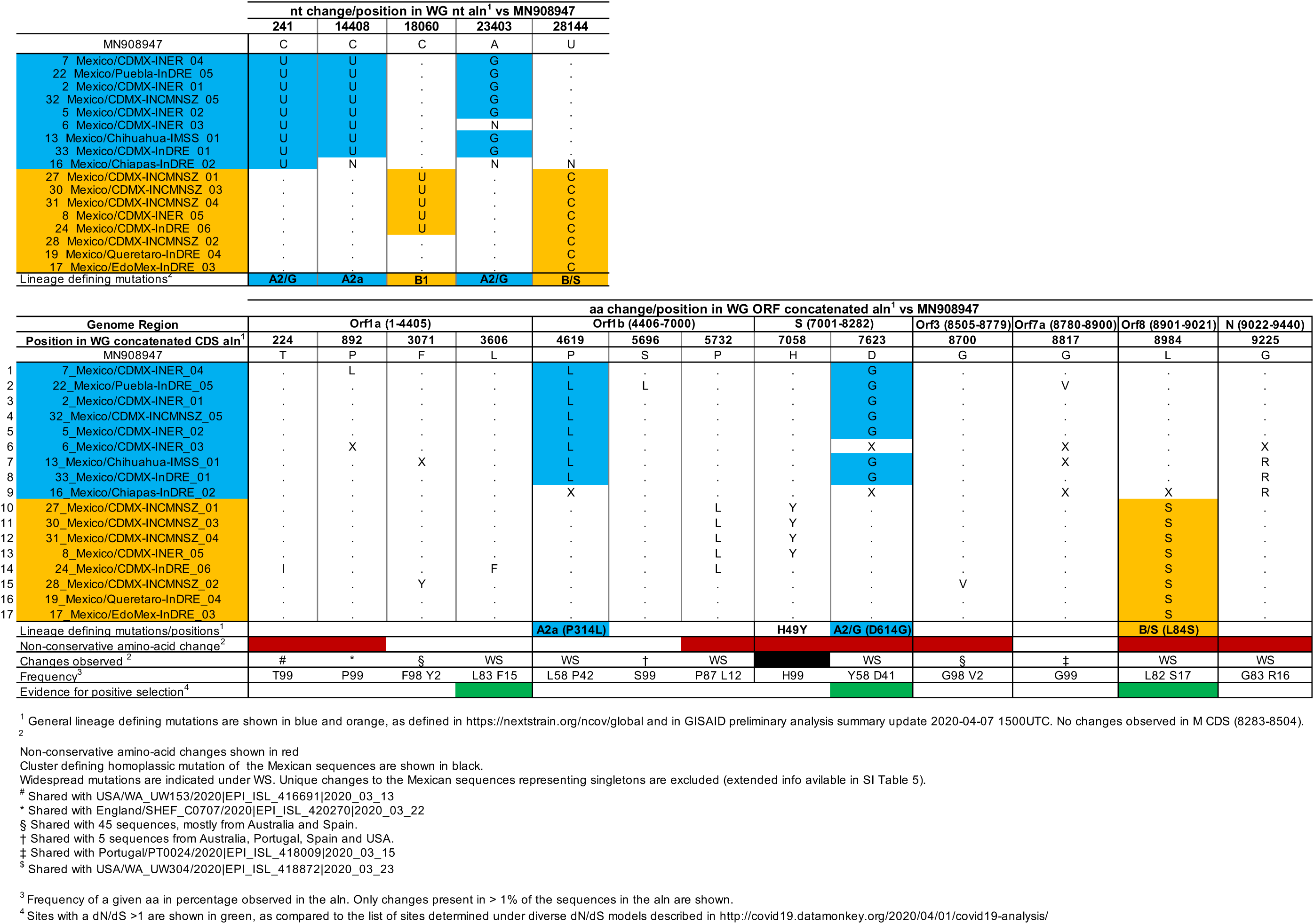
Nucleotide and amino acid changes in the Mexican samples compared to the reference strain Wuhan-Hu-1.

To visualize details on the phylogenetic relatedness of the Mexican sequences, we then subsampled the previous large-scale WG alignment in a phylogenetically-informed manner, as based on the position of the Mexican strains within the large-scale trees and by selecting sequences using pairwise genetic distances^25,26^. Briefly, all Mexican sequences were retained together with their immediate ancestors and descendants, and 184 sequences were further selected based on the minor pairwise genetic distance in relation to the Mexican genomes under a threshold of 99.5% (SI Table 3). The subsampled WG alignment was scanned for recombinant sequences using the GARD algorithm^27^ in the Datamonkey server^28,29^. A total of 201 sequences, including the Wuhan-Hu-1 reference genome, were used to re-estimate subsampled trees, as described above.

Global phylogenetic analysis and shared nucleotide substitution patterns^23^ were used to confirm the position of the Mexican sequences based on the consensus clustering patterns observed within the large-scale WG tree, the subsampled WG tree, and the individual large-scale ORF trees, using a bootstrap value >50 for branch support, when possible (SI Table 2). In general, we observed consistency within the global tree and our large-scale and subsampled trees, as depicted by a conserved general structure at an internal branch level. Finally, analysis of the phylogenetic relationship between the Mexican and other viral sequences to identify groups of introduction events (IE) and local transmissions (LT) was done based on the following local definition, in which and IE or LT must include: *i)* one or more Mexican sequences, *ii)* a minimum of one closest related sister sequence(s), *iii)* and/or the immediate common ancestor ^30,31^.

#### Mutation identification

Snippy^32^ was used to identify all mutations unique to the Mexican genomes, as compared to the reference genome sequence of isolate Wuhan-Hu-1. The large-scale WG alignment in nucleotides (including UTR and intergenic regions) was used as input for SI Table 4, while the large-scale WG concatenated ORF alignment was used as input for SI Table 5. The frequency and distribution for nucleotide and amino acid changes were determined using a normalized Sequence Logo, implemented by JalView^33^. Non-conservative amino acid changes were determined based on amino acid properties. Finally, the list of amino acid changes obtained was compared to the list of known sites scored to be evolving under pervasive, episodic or directional positive selection, as tested under several *dN/dS* models^34,35^.

## RESULTS AND DISCUSSION

### Multiple introduction events of SARS-CoV-2 variants from two different lineages

A total of 17 full viral genome sequences were obtained from selected Mexican samples representing the earliest sampled cases detected in the country (Figure 1). From the epidemiological data associated with the Mexican samples, 15 of the cases corresponded to introduction events from travelers returning from abroad that entered the country through Mexico City airport, with 5 of them then relocating to other places within the country using local transportation (either aerial or terrestrial). Two additional cases reported no travel history (Table 1). Global phylogenetic analysis^23^ confirmed that 8 Mexican strains (samples 8, 17, 19, 24, 27, 28, 30, 31) grouped within the SARS-CoV-2 lineage B (also called lineage S, composed by sequences predominantly from the Americas). The remaining 9 Mexican strains (samples 2, 5, 6, 7, 13, 16, 22, 32, and 33) grouped within lineage A2 and subclade A2a (also called lineage G, composed by sequences predominantly from Europe) (Figure 2)^36^. Both the A2/G and B/S lineages are contemporary, as the estimated date of emergence for lineage B/S is 12/29/2019 (CI: 12/20/2019, 01/03/2020), while for lineage A2/G is 01/18/2020 (CI 01/02/2020, 01/19/2020)^23^. The collection dates for the Mexican samples that fall within lineage B1 range from 03/04/2020 to 03/15/2020, while those that fall within lineage A2 range from 02/27/2020 to 03/15/2020, including the strain corresponding to the first reported case in México^37^ REF (sample 33). This suggests an initial co-circulation of both the A2/G and B/S lineages in Mexico (Figure 2). Viruses belonging to the third lineage reported, V, were not identified in this study.

**Figure 1.**
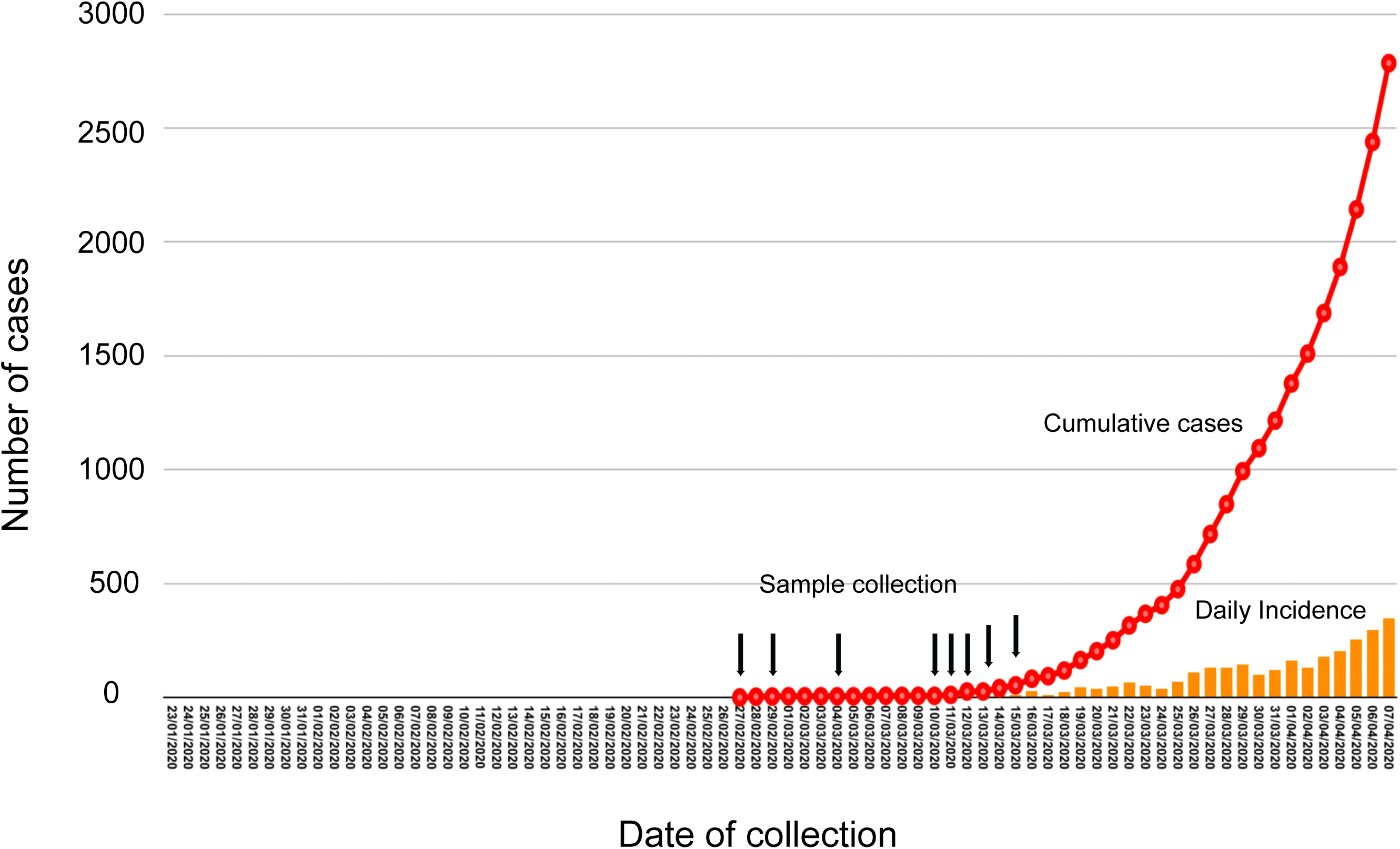
Epidemiological positioning of the SARS-CoV-2 samples from Mexico. Epidemiological curve for the early SARS-CoV-2 epidemic in Mexico, dating from late February until early April. The rise in cumulative cases is shown in red, whilst the daily incidence is shown with orange bars. Dates of collection for the samples used in this study are indicted with the black arrows. One arrow might indicate the collection of more than two samples (dates shown in Table 1).

**Figure 2.**
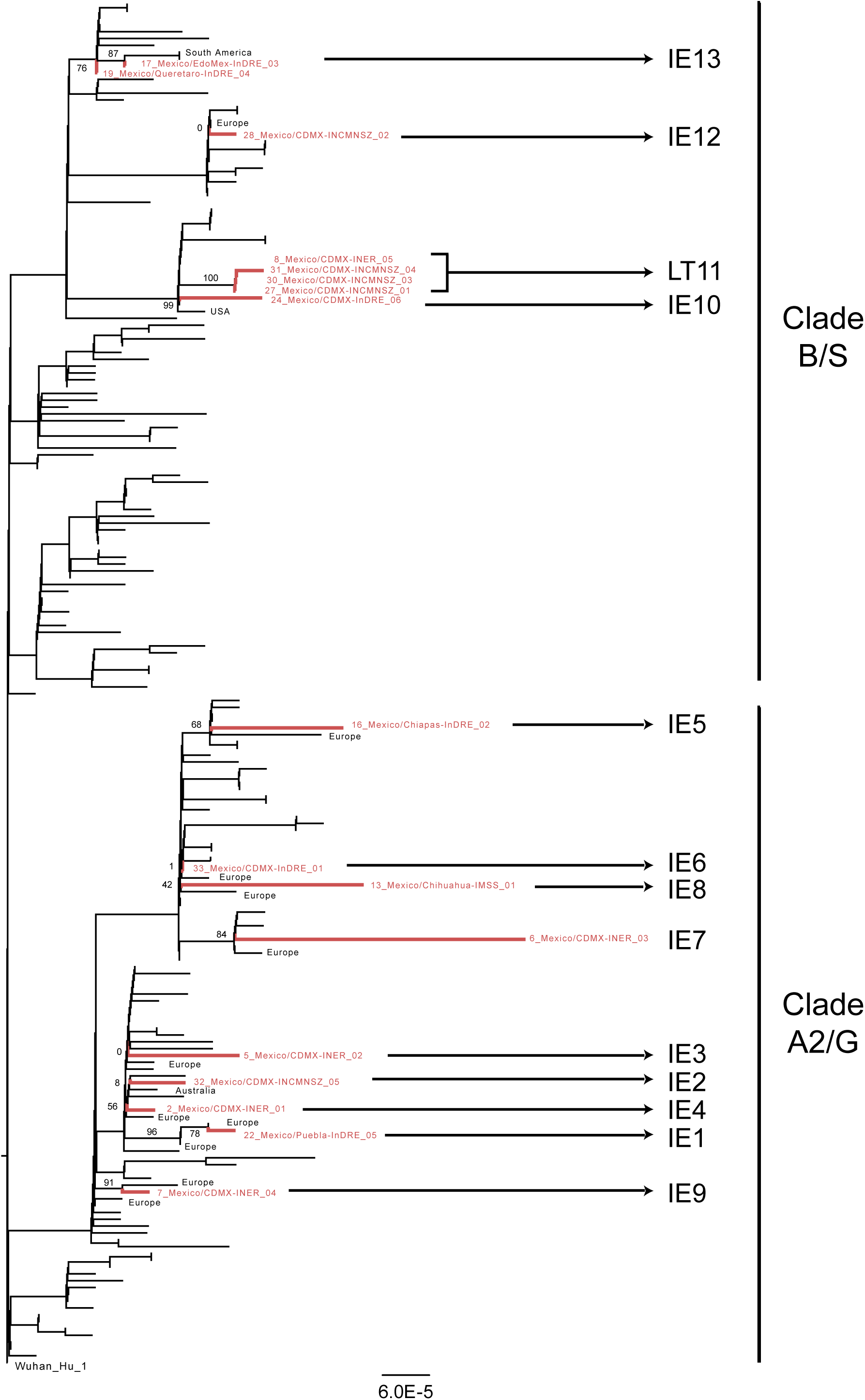
Phylogenomic positioning of the SARS-CoV-2 strains from Mexico. RAxML tree estimated from the reduced whole genome alignment built using the concatenated main viral ORFs, and rooted using the Wuhan-Hu-01 strain. Sequences corresponding to the viral isolates from Mexico sequenced in this work are shown in red, while the region of origin for the identified closest related immediate ancestors and/or sister isolates is indicated in black. Support values for the branches of interest are indicated with numbers next to the branches. The clusters identified in this work representing introduction events (IE1-IE10 and IE12 and IE13) and the local transmission (LT11) are shown next to the strain names.

Consensus clustering patterns observed within the large-scale and subsampled trees (SI Table 2), showed eight well-supported independent introduction events (IE1, IE3, IE5, IE7, IE9, IE10, IE12 and IE13; Figure 2). For IE2, IE4, IE6 and IE8, we were unable to determine their origins and the immediate phylogenetic relatedness of the Mexican sequences, due to low support values and inconsistent clustering patterns across trees (SI Table 2). The current resolution of phylogenomic analyses is limited by the low diversity of the SARS-CoV-2 virus. Thus, for the characterized Mexican viral strains, we could only determine with confidence the geographical origins at a regional level, rather than at country-level resolution. Altogether, these observations suggest that the virus variants identified in this study were more closely related to virus strains circulating at the time in the USA and Europe rather than in China or South East Asia.

### Evidence for early local transmission

Sequences 27 and 31 correspond to two individuals that shared travel history to Vail, CO, USA, and that were in direct contact with case 28 in the return flight to Mexico. Nonetheless, the distribution of sequence 28 in an independent group (IE12) in relation to sequences 27 and 31 (Figure 2, SI Table 2), suggests that there were at least two different viral variants co-circulating within this specific location in the USA. On the other hand, sequences 8, 27, 30 and 31 grouped together, representing a local transmission cluster (LT11) (Figure 2). This observation is supported by phylogenetic consistency within our local trees and the global tree^23^, and a high support value (bs: 100) observed for LT11 in all cases (S1 Table 2, Figure 2). Case 8 corresponds to an individual with no epidemiological relationship or contact with cases 27, 30 and 31, and with no travel history. Case 30 had travel history to Madrid, Spain, but no direct exposure with cases 27 and 31. For case 30, the possibility of this person acquiring the virus whilst being abroad cannot be ruled out. However, our analysis shows evidence that this person may have contracted the virus whilst already being in Mexico. Thus, LT11 strongly supports the occurrence of at least one independent local transmission event in Mexico City (case 8), as early as the second week of March.

### Genetic variation within the Mexican viral genomes

When compared to the Wuhan-Hu-1 reference genome, the Mexican sequences displayed between 4 and 10 nucleotide substitutions, and between 1 and 5 amino acid changes. This is consistent with the reported rate of evolution of ∼8×10^−4^ nucleotide substitutions per site per year, equivalent to ∼2 substitutions per month^38,39^. Collectively, 46 nucleotide substitutions and 20 amino acid residues were identified within the Mexican viral genomes (SI Table 4 and 5). As expected, the majority of these variants were not conserved through the genomes, and only 15 nucleotide changes and 6 amino acid changes were shared by more than one sequence. These results exclude sequences 6, 13 and 16, which showed a considerably lower coverage and depth when compared to the remaining 14 high-quality viral genomes obtained. Thus, most of the variability observed could be explained by errors introduced during reverse transcription, PCR amplification, sequencing or assembly (SI Table 1 and 4).

Consistent with our phylogenomic analysis, all Mexican sequences belonging to lineage A2 showed two clade-specific nucleotide substitutions, C241T and A23403G. A23403G results in the D614Ge amino acid change in the Spike protein (Table 2). All lineage A2a sequences (including the Mexican isolates) had the additional nucleotide substitution C14408T, resulting in the amino acid change P314L in Orf1b^23^. Sequences within lineage B had the nucleotide substitution T28144C rendering the L84S amino acid change in the Orf8 protein. Similarly, lineage B1 sequences also showed the nucleotide substitution C18060T (Table 2). No evidence for recombination was found within any of the alignments, in agreement with previous observations^28^. Taken together, our results suggest the Mexican viral sequences display the genetic changes according to their phylogenetic placement.

### H49Y amino acid change in Mexican sequences within LT1

We further identified within all Mexican sequences grouping with the local transmission cluster (LT11), an H to Y amino acid change in position 49 of the Spike protein, corresponding to the C21707T nucleotide change in the whole genome and nucleotide alignment numbering. H49Y represents a non-conservative mutation located within the N-terminal domain (NTD) of the S glycoprotein trimer^40^. No evidence of episodic or directional positive selection was found for this site, as tested under several *dN/dS* models^34,35^.

Mapping this nucleotide change onto the Nextstrain global tree^23^ with dates ranging from December 2019 to April 2020, revealed that this mutation represents a homoplasy that has occurred with a frequency of 0.4% (20/4533) within the genomes displayed in the global tree as of May 6^th^ 2020. This change first occurred in a single cluster of 14 sequences mainly from China (representative sequence: Jiangsu/JS02/2020 EPI ISL 411952). The estimated date of emergence for this cluster that eventually died off (stopped circulating and had no more descendants) was of 01/12/2020 (CI:01/08/2020-01/16/2020). This mutation was detected again in the Mexican sequences in LT11 (Table 1, Figure 2). Since then, the C21707T/H49Y mutation has also appeared independently in three strains from Australia (EPI ISL 426898, EPI_ISL_430632 and EPI_ISL_430633), two strains from the UK (EPI_ISL_433757 and EPI_ISL_432818), one strain from Taiwan (EPI_ISL_417518) and two strains from the USA (EPI_ISL_408010 and EPI_ISL_430050), with collection dates ranging from 01/29/2020 to 04/05/2020 (sequences deposited on GISAID platform by May 10^th^ 2020). Of notice, all the sequences displaying the H49Y mutation belong to different viral lineages (either, A2, B1 or others), confirming non-phylogenetic correlation (e.g. founder effect), but rather an independent occurrence. It would be important to follow up the genomic surveillance of homoplasic mutations that keep emerging independently through time and space, either to determine if these are sequencing errors^41^, or to know if these are real homoplasies that may be associated with changes in biological properties or the virus, such as increased virulence as has been shown for other viruses^25^. In addition, using H49Y as a molecular marker could be used to follow up any further circulation and spread of these viral variants in Mexico.

## Data Availability

The generated sequences SARS-CoV-2 from Mexico can be found in GISAID and in Nextstrain^24^. The corresponding GISAID accession numbers are listed in Table 1.

## ACKNOWLEDGEMENTS

This work was partially supported by grant “Epidemiología genómica de los virus SARS-CoV-2 circulantes en México” from the National Council for Science and Technology (CONACyT)-Mexico to CFA, and by grants from the Mexican Government (Comisión de Equidad y Género de las Legislaturas LX-LXI y Comisión de Igualdad de Género de la Legislatura LXII de la H. Cámara de Diputados de la República Mexicana) to SAR and CB. MEZ is supported by a Leverhulme Trust ECR Fellowship (ECF-2019-542). We thank the “Unidad de Secuenciación Masiva y Bioinformática” of the “Laboratorio Nacional de Apoyo Tecnológico a las Ciencias Genómicas” (CONACyT #260481) for their support in sequencing services. We thank all the staff of the Technological Development and Molecular Research Unit, Virology Department, and Sample Control and Services Department at InDRE for their technical assistance. The findings and conclusions in this report are only the responsibility of the authors and do not necessarily of the institutions involved.

---

**Supplementary Table 1.**
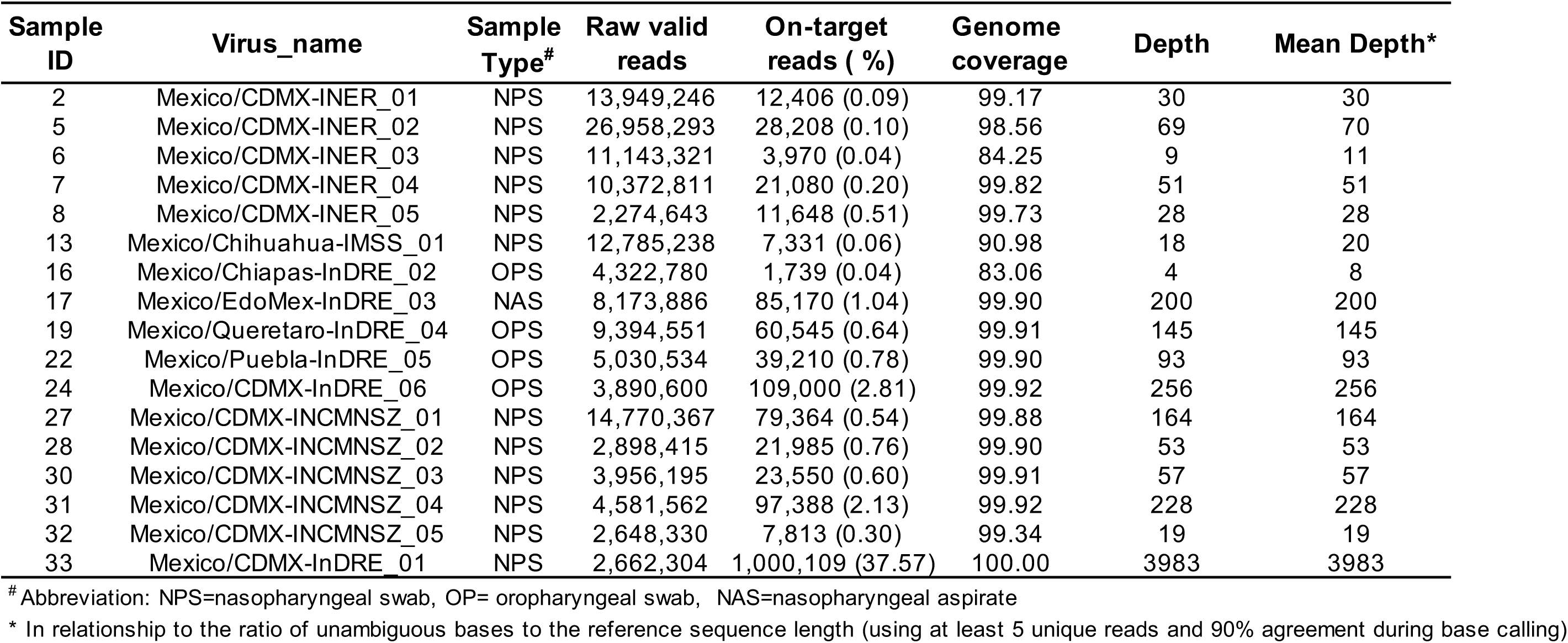
Sequencing results.

| Sample ID | Virus_name | Sample Type<sup>#</sup> | Raw valid reads | On-target reads ( %) | Genome coverage | Depth | Mean Depth<sup>*</sup> |
| --- | --- | --- | --- | --- | --- | --- | --- |
| 2 | Mexico/CDMX-INER_01 | NPS | 13,949,246 | 12,406 (0.09) | 99.17 | 30 | 30 |
| 5 | Mexico/CDMX-INER_02 | NPS | 26,958,293 | 28,208 (0.10) | 98.56 | 69 | 70 |
| 6 | Mexico/CDMX-INER_03 | NPS | 11,143,321 | 3,970 (0.04) | 84.25 | 9 | 11 |
| 7 | Mexico/CDMX-INER_04 | NPS | 10,372,811 | 21,080 (0.20) | 99.82 | 51 | 51 |
| 8 | Mexico/CDMX-INER_05 | NPS | 2,274,643 | 11,648 (0.51) | 99.73 | 28 | 28 |
| 13 | Mexico/Chihuahua-IMSS_01 | NPS | 12,785,238 | 7,331 (0.06) | 90.98 | 18 | 20 |
| 16 | Mexico/Chiapas-InDRE_02 | OPS | 4,322,780 | 1,739 (0.04) | 83.06 | 4 | 8 |
| 17 | Mexico/EdoMex-InDRE_03 | NAS | 8,173,886 | 85,170 (1.04) | 99.90 | 200 | 200 |
| 19 | Mexico/Queretaro-InDRE_04 | OPS | 9,394,551 | 60,545 (0.64) | 99.91 | 145 | 145 |
| 22 | Mexico/Puebla-InDRE_05 | OPS | 5,030,534 | 39,210 (0.78) | 99.90 | 93 | 93 |
| 24 | Mexico/CDMX-InDRE_06 | OPS | 3,890,600 | 109,000 (2.81) | 99.92 | 256 | 256 |
| 27 | Mexico/CDMX-INCMNSZ_01 | NPS | 14,770,367 | 79,364 (0.54) | 99.88 | 164 | 164 |
| 28 | Mexico/CDMX-INCMNSZ_02 | NPS | 2,898,415 | 21,985 (0.76) | 99.90 | 53 | 53 |
| 30 | Mexico/CDMX-INCMNSZ_03 | NPS | 3,956,195 | 23,550 (0.60) | 99.91 | 57 | 57 |
| 31 | Mexico/CDMX-INCMNSZ_04 | NPS | 4,581,562 | 97,388 (2.13) | 99.92 | 228 | 228 |
| 32 | Mexico/CDMX-INCMNSZ_05 | NPS | 2,648,330 | 7,813 (0.30) | 99.34 | 19 | 19 |
| 33 | Mexico/CDMX-InDRE_01 | NPS | 2,662,304 | 1,000,109 (37.57) | 100.00 | 3983 | 3983 |

<sup>#</sup> Abbreviation: NPS=nasopharyngeal swab, OP= oropharyngeal swab, NAS=nasopharyngeal aspirate
<sup>\*</sup> In relationship to the ratio of unambiguous bases to the reference sequence length (using at least 5 unique reads and 90% agreement during base calling)

**Supplementary Table 2.** Clustering patterns for the Mexican samples within the reduced and large-scale WG and individual ORF ML trees.

| reduced -WG<sup>1</sup> | large-scale WG<sup>1</sup> | large-scale Orf1a<sup>1</sup> | large-scale Orf1b<sup>1</sup> | large-scale S<sup>1</sup> |
| --- | --- | --- | --- | --- |
| IE1-bs 78<br>22_Mexico/Puebla-InDRE_05/2020\|EPI_ISL_424672\|Mexico_Puebla\|31\|M\|EH_Spain_France\|3_11_2020<br>Portugal/PT0024/2020\|EPI_ISL_418009\|PRT_Portugal\|Age_29\|Sex_F\|D0_NA\|EH_NA\|3_15_2020<br>Basal-bs 96: Spain/Madrid_H11_40/2020\|EPI_ISL_417967\|ESP_Madrid\|Age_44\|Sex_F\|D0_NA\|EH_NA\|3_12_2020 | IE1-bs 6<br>YES | IE1-bs 0<br>NO | IE1-bs 0<br>YES | IE1-bs 0<br>NO-POLYTOMY |
| IE2-bs 8<br>32_Mexico/CDMX-INCMNSZ_05/2020\|EPI_ISL_426365\|Mexico_CDMX\|70\|F\|EH_Vail_USA\|3_12_2020<br>Australia/VIC254/2020\|EPI_ISL_419949\|AUS_Victoria\|Age_67\|Sex_M\|D0_NA\|EH_NA\|2020_03_21<br>Basal-bs 0: Australia/VIC164/2020\|EPI_ISL_419861\|AUS_Victoria\|Age_37\|Sex_NA\|D0_NA\|EH_NA\|2020_03_18 | IE2-bs 27<br>YES | IE2-bs 0<br>NO | IE2-bs 0<br>NO | IE2-bs 0<br>NO |
| IE3-bs 0<br>5_Mexico/CDMX-INER_02/2020\|EPI_ISL_424348\|Mexico_CDMX\|55\|F\|EH_Spain_France\|3_13_2020<br>Spain/Madrid_H12_1905/2020\|EPI_ISL_421175\|ESP_Madrid\|Age_25\|Sex_F\|D0_NA\|EH_NA\|2020_03_29<br>Basal-bs: None | IE3-bs 0<br>YES | IE3-bs 0<br>NO | IE3-bs 0<br>YES-POLYTOMY | IE3-bs 0<br>NO |
| IE4-bs 56<br>2_Mexico/CDMX-INER_01/2020\|EPI_ISL_424345\|Mexico_CDMX\|42\|M\|EH_Spain\|3_12_2020<br>Hungary/2/2020\|EPI_ISL_418183\|HUN_Baranya\|Age_26\|Sex_F\|D0_H\|EH_NA\|2020_03_17<br>Basal-bs 0: None | IE4-bs 61<br>NO | IE4-bs 0<br>NO | IE4-bs 0<br>NO-POLYTOMY | IE4-bs 0<br>NO |
| IE5-bs 68<br>Wales/PHWC_25551/2020\|EPI_ISL_421004\|GBR_Wales\|Age_73\|Sex_M\|D0_NA\|EH_NA\|2020_03_23<br>16_Mexico/Chiapas-InDRE_02/2020\|EPI_ISL_424666\|Mexico_Chiapas\|18\|F\|EH_Italy\|2_29_2020<br>Basal-bs 0: None | IE5-bs 19<br>YES | IE5-bs 0<br>NO | IE5-bs 0<br>NO | IE5-bs 0<br>NO-POLYTOMY |
| IE6-bs 0<br>33_Mexico/CDMX-InDRE_01/2020\|EPI_ISL_412972\|Mexico_CDMX\|35\|M\|EH_Italy\|2_27_2020<br>Basal-bs 1: England/20122119502/2020\|EPI_ISL_418681\|GBR_England\|Age_31\|Sex_F\|D0_NA\|EH_NA\|2020_03_17 | IE6-bs 1<br>NO | IE6-bs 0<br>NO-POLYTOMY | IE6-bs 0<br>NO-POLYTOMY | IE6-bs 0<br>NO-POLYTOMY |
| IE7-bs 84<br>6_Mexico/CDMX-INER_03/2020\|EPI_ISL_424625\|Mexico_CDMX\|29\|M\|EH_Spain\|3_13_2020<br>England/20108034006/2020\|EPI_ISL_417258\|GBR_England\|Age_74\|Sex_F\|D0_NA\|EH_NA\|2020_03_06 | IE7-bs 67<br>YES | IE7-bs 0<br>YES | IE7-bs 27<br>YES | IE7-bs 0<br>NO |
| IE8-bs 42<br>Italy/TE4959/2020\|EPI_ISL_418259\|ITA_Abruzzo\|Age_76\|Sex_M\|D0_NA\|EH_NA\|2020_03_14<br>13_Mexico/Chihuahua-IMSS_01/2020\|EPI_ISL_424731\|Mexico_Chihuahua\|21\|M\|EH_UK_France_USA\|3_13_2020<br>Basal-bs 0: None | IE8-bs 18<br>YES | IE8-bs 0<br>NO | IE8-bs 0<br>NO-POLYTOMY | IE8-bs 0<br>NO-POLYTOMY |
| IE9-bs 91<br>7_Mexico/CDMX-INER_04/2020\|EPI_ISL_424626\|Mexico_CDMX\|25\|F\|EH_Egypt_Barcelona_Spain\|3_15_2020<br>England/SHEF_C0707/2020\|EPI_ISL_420270\|GBR_England\|Age_73\|Sex_F\|D0_NA\|EH_NA\|2020_03_22<br>Basal-bs: None | IE9-bs 79<br>YES | IE9-bs 0<br>YES | IE9-bs 0<br>NO-POLYTOMY | IE9-bs 0<br>NO |
| IE10-bs 99<br>Australia/VIC110/2020\|EPI_ISL_419716\|AUS_Victoria\|Age_62\|Sex_F\|D0_NA\|EH_NA\|2020_03_18<br>24_Mexico/CDMX-InDRE_06/2020\|EPI_ISL_424673\|Mexico_CDMX\|63\|F\|EH_Denver_USA\|3_12_2020 | IE10-bs 0<br>NO | IE10-bs 2<br>YES | IE10-bs 0<br>YES-POLYTOMY | IE10-bs 0<br>YES-POLYTOMY |
| LT11-bs 100 CLUSTER<br>31_Mexico/CDMX-INCMNSZ_04/2020\|EPI_ISL_426364\|Mexico_CDMX\|70\|M\|EH_Vail_USA\|3_12_2020<br>8_Mexico/CDMX-INER_05/2020\|EPI_ISL_424627\|Mexico_CDMX\|38\|M\|EH_None\|3_15_2020<br>27_Mexico/CDMX-INCMNSZ_01/2020\|EPI_ISL_426361\|Mexico_CDMX\|32\|M\|EH_None\|3_12_2020<br>30_Mexico/CDMX-INCMNSZ_03/2020\|EPI_ISL_426363\|Mexico_CDMX\|38\|M\|EH_None\|3_12_2020<br>Basal-bs: None | IE11-bs 98 CLUSTER<br>YES | C11.1-bs 0<br>BROKEN CLUSTER | C11.1-bs 0<br>BROKEN CLUSTER | IE11-bs 89 CLUSTER<br>YES |
| IE12-bs 2<br>28_Mexico/CDMX-INCMNSZ_02/2020\|EPI_ISL_426362\|Mexico_CDMX\|71\|M\|EH_Vail_USA_Germany\|3_10_2020<br>Basal-bs 0: Spain/CastillaLeon201372/2020\|EPI_ISL_418249\|ESP_Castilla_y_Leon\|Age_25\|Sex_F\|D0_NA\|EH_NA\|2020_03_03 | IE12-bs 0<br>YES | IE12-bs 0<br>YES | IE12-bs 0<br>YES-POLYTOMY | IE12-bs 0<br>NO |
| IE13 -bs 87<br>Chile/Talca_2/2020\|EPI_ISL_414578\|CHL_Talca\|Age_33\|Sex_F\|D0_NA\|EH_NA\|2020_03_04<br>Chile/Talca_1/2020\|EPI_ISL_414577\|CHL_Talca\|Age_33\|Sex_M\|D0_R\|EH_Asia_Europe\|2020_03_02<br>17_Mexico/EdoMex-InDRE_03/2020\|EPI_ISL_424667\|Mexico_Edomex\|71\|M\|EH_Italy\|3_4_2020<br>Basal-bs 2: 19_Mexico/Queretaro-InDRE_04/2020\|EPI_ISL_424670\|Mexico_Queretaro\|43\|M\|EH_Spain_Holand_USA\|13_10_2020 | IE13-bs 0<br>YES | IE13-bs 0<br>YES | IE13-bs 33<br>YES | IE13-bs 0<br>NO |
|  | NO-POLYTOMY-BROKEN CLUSTER WITH 19 | NO-POLYTOMY-BROKEN CLUSTER WITH 19 | NO-POLYTOMY-BROKEN CLUSTER WITH 19 | NO |

<sup>1</sup>Consistency for clustering patterns observed within >2 trees and/or bootstrap support values >50 are indicated in red. YES/NO indicates conserved clustering patterns.

**Supplementary Table 3.**
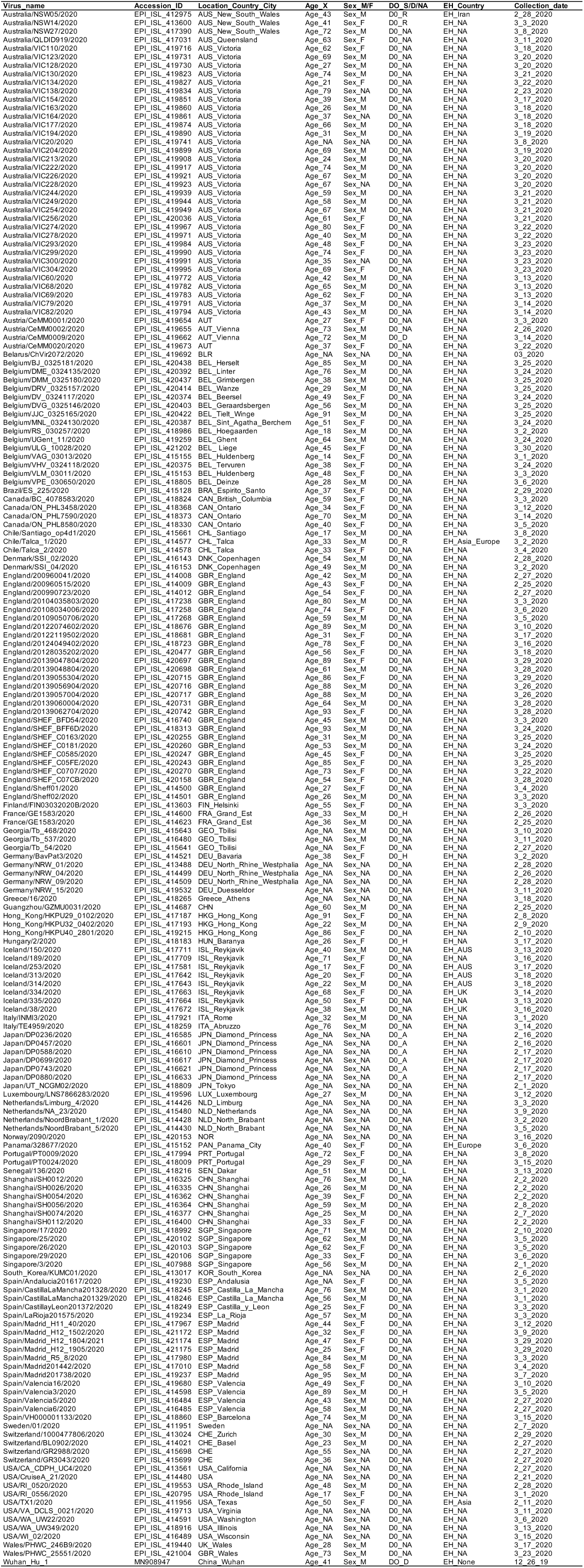
List of sequences and corresponding metadata used for the reduced alignments.

**Supplementary Table 4.**
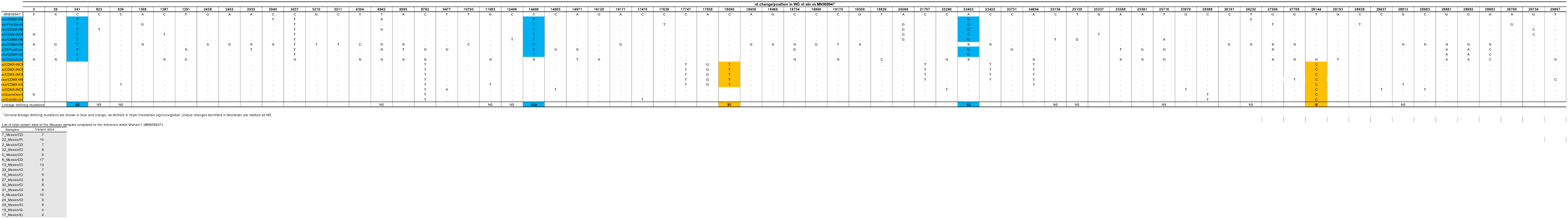
Nucleotide changes within the Mexican samples compared to the reference strain Wuhan-Hu-1. Sequences were order by their philogenetic position in Figure 2.

**Supplementary Table 5.**
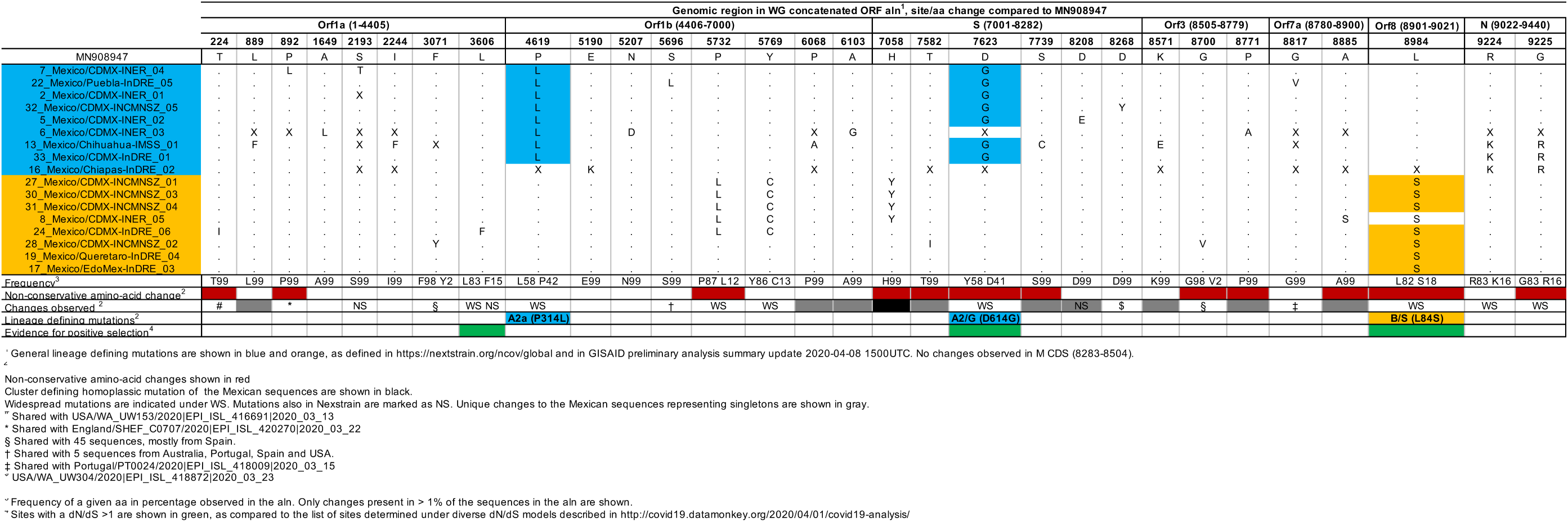
Amino acid changes within the Mexican samples compared to the reference strain Wuhan-Hu-1.

